# Microbially-conjugated Bile Salts Found in Human Bile Activate the Bile Salt Receptors TGR5 and FXR

**DOI:** 10.1101/2023.09.04.556292

**Authors:** Ümran Ay, Martin Leníček, Raphael S. Haider, Arno Classen, Hans van Eijk, Kiran V.K. Koelfat, Gregory van der Kroft, Ulf. P. Neumann, Carsten Hoffmann, Carsten Bolm, Steven W.M. Olde Damink, Frank G. Schaap

**Affiliations:** Department of General, Visceral and Transplant Surgery, University Hospital Aachen, 52074 Aachen, Germany; Institute of Medical Biochemistry and Laboratory Diagnostics, Faculty General Hospital and 1^st^ Faculty of Medicine, Charles University, 12808 Prague, Czech Republic; Institute of Molecular Cell Biology, Center for Molecular Biomedicine, Jena University Hospital, Jena, Germany; Division of Physiology, Pharmacology and Neuroscience, School of Life Sciences, Queen’s Medical Center, University of Nottingham, Nottingham, United Kingdom; Center of Membrane Protein and Receptors, Universities of Birmingham and Nottingham, Midlands, United Kingdom; Institute of Organic Chemistry, RWTH Aachen University, 52074 Aachen, Germany; Department of Surgery, NUTRIM School of Nutrition and Translational Research in Metabolism, Maastricht University, 6200 MD Maastricht, The Netherlands

## Abstract

**Background & Aims:** Bile salts of hepatic and microbial origin mediate inter-organ crosstalk in the gut-liver axis. Here, we assessed whether the newly discovered class of microbial bile salt conjugates (MBSCs), activate the main host bile salt receptors (TGR5 and FXR) and enter the human systemic and enterohepatic circulation.

**Approach & Results:** *N*-amidates of (chenodeoxy)cholic acid and leucine, tyrosine and phenylalanine were synthesized. Receptor activation was studied in cell-free and cell-based assays. MBSCs were quantified in mesenteric and portal blood and bile of patients undergoing pancreatic surgery. MBSCs were activating ligands of TGR5 as evidenced by recruitment of G_sα_ protein, activation of a cAMP-driven reporter, and diminution of LPS-induced cytokine release from macrophages. Intestine- and liver-enriched FXR isoforms were both activated by MBSCs, provided that a bile salt importer was present. Affinity of MBSCs for TGR5 and FXR was not superior to host-derived bile salt conjugates. Individual MBSCs were generally not detected (*i*.*e*. <2.5 nmol/L) in human mesenteric or portal blood, but Leu- and Phe-variants were readily measurable in bile, where MBSCs comprised up to 213 ppm of biliary bile salts.

**Conclusions:** MBSCs activate the cell surface receptor TGR5 and the transcription factor FXR, and are substrates for intestinal (ASBT) and hepatic (NTCP) transporters. Their entry into the human circulation is, however, non-substantial. Given low systemic levels and surplus of other equipotent bile salt species, the studied MBSCs are unlikely to have an impact on enterohepatic TGR5/FXR signaling in humans. Origin and function of biliary MBSCs remain to be determined.

**Graphical Abstract:** 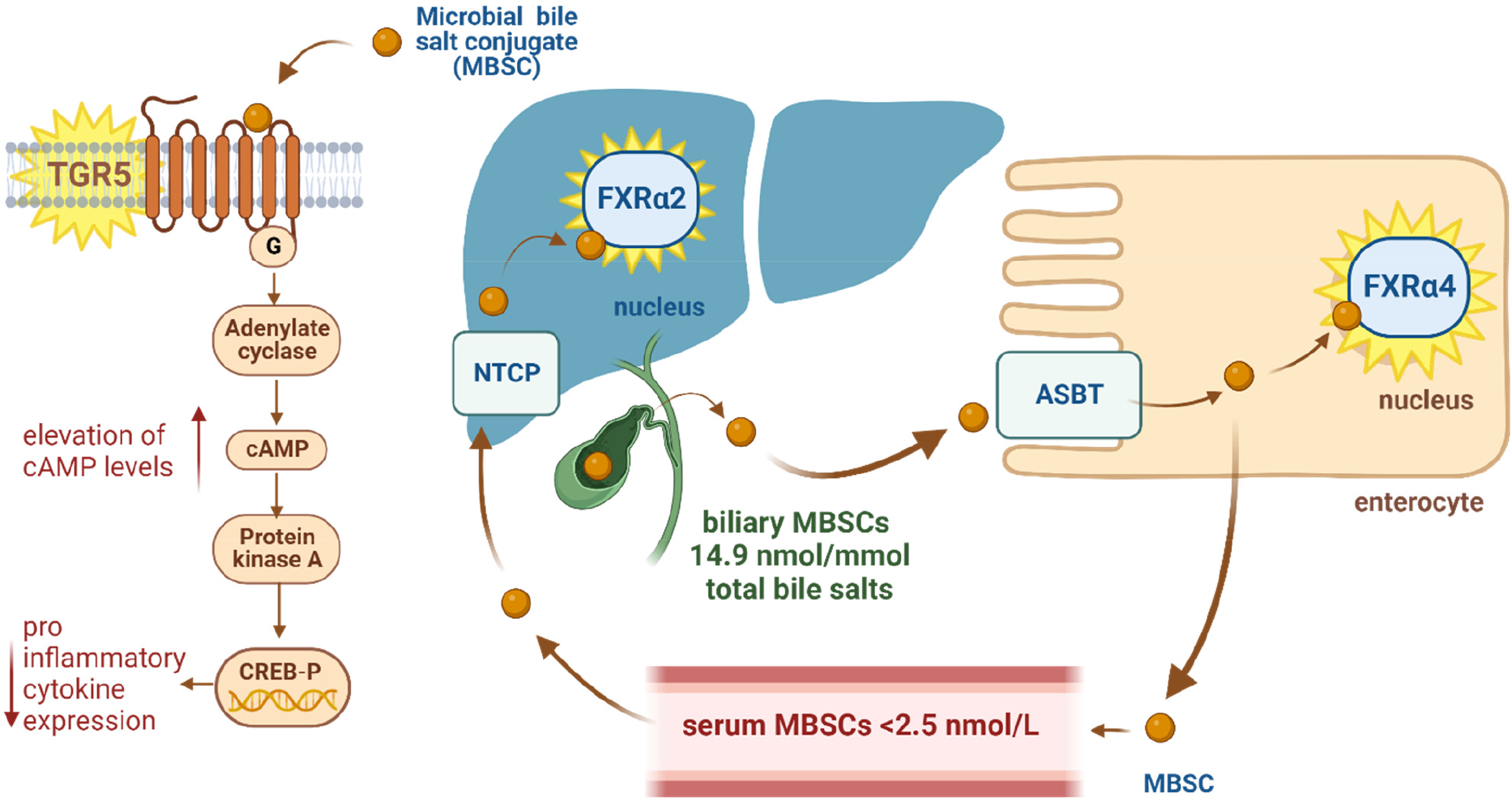

Created with BioRender.com

## Introduction

The liver plays a central role in the handling of bile salts, a class of cholesterol-derived, amphipathic molecules with digestive and signaling actions (1). Bile salt homeostasis is deranged in numerous liver diseases, and direct pathogenic contributions of bile salts are evident in cholestatic liver disorders (2-5). Bile salts are molecular mediators of inter-organ crosstalk in the gut-liver axis, whose proper function is key to maintaining intestinal and liver health (6,7).

The microbial community in the gut has the unique ability to transform liver-derived (*i*.*e*. primary) bile salts during their enterohepatic cycle. Microbial conversion of bile salts appears confined to bacteria, with a range of gram-negative and gram-positive species harboring one or multiple enzymes that act on the side chain conjugate group or at specific positions within the steroid nucleus (8). Some of these secondary bile salts, notably deoxycholic acid (DCA) and conjugates thereof, can become a substantial part of the recirculating pool, impacting host physiology by affecting the digestive, bactericidal and signaling properties of bile salts (8). Less prominent secondary species may act more locally, *e*.*g*. close to the site of formation, as appears the case for newly recognized bile salt metabolites that modulate T cell differentiation in the intestines (9-11).

A new class of bacterial bile salt metabolites was first reported in 2020 (12), and comprises C_24_ bile salts that are *N*-amidated with amino acids other than those used by the host conjugation system (*i*.*e*. glycine and taurine) (13). These microbial bile salt conjugates (MBSCs) were present in human and mouse feces, could not be detected in germ-free mice, and were formed *in vitro* by *Clostridium bolteae* after presenting with cholic acid (CA) and amino acids. The initial mass spectrometry-based approach led to identification of three MBSCs, *viz*. leucocholic acid (LeuCA), phenylalanocholic acid (PheCA) and tyrosocholic acid (TyrCA), but this number rapidly expanded to >100 by targeted efforts that revealed broad diversity at the level of both bile acid and amino acid precursor (14-16). Moreover, numerous additional bacterial producers were identified (14,17), and led to the notion that MBSC formation is common among human gut bacteria. Amine *N*-acyl transferase activity of bile salt hydrolase (*bsh*), a bacterial enzyme known to catalyze bile salt deconjugation, has been implicated in the generation of MBSCs. Additional biosynthetic routes likely exist as bacteria devoid of *bsh* can form MBSCs (14).

The biological and pathological functions of MBSCs remain enigmatic (18, 19). MBSCs likely act as ligands for host bile salt receptors including Farnesoid X Receptor (FXR) and Pregnane X Receptor (PXR), and accordingly have potential to modulate processes controlled by these nuclear receptors (18,20). A recent study demonstrated that several MBSCs inhibit germination of *C. difficile* spores *in vitro*, pointing towards a role of MBSCs in bacterial communication (14). Analysis of public mass spectrometry data sets revealed an association between fecal MBSC abundance (*viz*. LeuCA, PheCA, TyrCA) and inflammatory bowel disease, notably Crohn’s disease with dysbiosis (12). Mice given a high-fat diet had higher abundance of aforementioned MBSCs in their feces, hinting at a relation with obesity-related disorders (12).

Quantitative data on levels in biospecimens other than intestinal content is lacking, and it is unresolved if MBSCs enter the host circulation. Of note, repeated oral gavage of mice with a considerable dose of LeuCA or TyrCA, did not lead to detection of these MBSCs in portal blood or bile (12). We previously reported that MBSCs comprise a minor fraction (up to 40 ppm) of total bile salts in chyme of patients with intestinal failure (18). Clearly, insight into affinities for bile salt receptors and quantitative data on levels in the circulation is pivotal to further our understanding of the role of MBSCs in health and disease. In this study we aimed to determine whether MBSCs are substrates for bile salt uptake transporters, study their interaction with the main bile salt receptors TGR5 and FXR, and assess their entry into the human circulation.

## Methods

### Materials

Chenodeoxycholic acid (CDCA), cholic acid (CA), obeticholic acid (OCA), taurolithocholic acid (TLCA), glycochenodeoxycholic acid (GCDCA), and Forskolin A were purchased from Sigma. Stocks were prepared in molecular biology grade DMSO (Sigma, D8418). MBSCs employed in this study were those initially discovered by Quinn *et al*. (12), *viz*. leucocholic acid (LeuCA), phenylalanocholic acid (PheCA) and tyrosocholic acid (TyrCA), as well as the respective *N*-amidates with CDCA (LeuCDCA, PheCDCA, TyrCDCA). Both *D*- and *L*-stereoisomers were used for *N*-amidation with Tyrosine. Chemical synthesis and purification of above MBSCs was essentially according to Quinn *et al*. (12), and all compounds were verified by ^1^H and ^13^C NMR in deuterated DMSO.

### Patient samples

Blood (superior mesenteric vein, inferior mesenteric vein, portal vein) and gallbladder bile samples were collected intraoperatively from patients undergoing pancreaticoduodenectomy for treatment of head of pancreas tumors at the University Hospital Aachen. These samples were collected in the framework of the FOCUS study (Ethical Commission University Hospital Aachen approval: EK 172/17), following written informed consent of the patient. Furthermore, plasma samples (systemic blood) were analyzed from a previously reported cohort of patients with acute intestinal failure undergoing chyme reinfusion (21). All research was conducted in accordance with both the Declarations of Helsinki and Istanbul.

### Cell culture

Human embryonic kidney cells (293T, ACC 635) were directly purchased from the German Collection of Microorganisms and Cell Cultures (DSMZ, Braunschweig, Germany). For the nanoBRET assay, 293 cells were directly acquired from the American Type Culture Collection (ATCC, CRL-1573). Murine macrophages (RAW264.7) were straight from ATCC (TIB-71). All cells were cultured in a humidified atmosphere containing 5% CO_2_ at 37°C in high-glucose DMEM supplemented with 10% fetal bovine serum, 1.0 mM pyruvate, 100 U/mL penicillin G and 100 μg/mL streptomycin. For serum-free conditions, the serum component was substituted for 0.2% BSA. All cell culture reagents were purchased from ThermoFisher Scientific.

In initial tests, cell viability was evaluated at the end of incubations by a dehydrogenase activity assay (Cell Counting Kit 8, Abcam #ab228554), and expressed relative to the solvent control.

### Plasmids

Expression vectors for human TGR5 (pCMV3-GPBAR1 C-GFPSpark, hereafter pCMV3-TGR5) and its control (pCMV3-C-GFPSpark, hereafter pCMV3-GFP), human ASBT (pCMV3-SLC10A2) and human NTCP (pCMV3-SLC10A1) were purchased from SinoBiological. Reporter plasmids were obtained from Promega and encompassed Firefly luciferase under the control of a cAMP-responsive element (pGL4.29[luc2P/CRE/Hygro], hereafter pGL4-CRE-Fluc) and *Renilla* luciferase (pRL-TK). An FXR reporter (pGL3-*Shp_e*-FLuc) and pcDNA3.1-based plasmids for expression of human FXRα2, FXRα4 and RXRα were kind gifts of Prof. Saskia van Mil (University Medical Center Utrecht, The Netherlands) and have been described elsewhere (22). Plasmids were propagated in TOP10 *E. coli* (Thermo Fisher Scientific) and endotoxin-free plasmid DNA was isolated using anion exchange columns (NucleoBond, Macherey-Nagel).

### NanoBRET assay

Ligand-induced recruitment of a mini-G_s_ protein to TGR5 was assessed by a nano-luciferase bioluminescence resonance energy transfer assay (nanoBRET) as detailed elsewhere (23, 24). In brief, 293 cells were transiently transfected (polyethylenimine, Sigma-Aldrich) with a mini-G_s_ protein and human TGR5 *C*-terminally fused with nano-luciferase, and seeded into white opaque 96-well plates the next day. Test compounds including solvent (0.5% DMSO) and positive controls (100 μM TLCA) were added 24 hours later, and donor and acceptor emissions were measured directly. BRET ratios (acceptor:donor emission) are expressed relative to the ratio of the solvent control (*i*.*e*. 0.5% DMSO = 1.0). Dose-response curves were fitted by non-linear regression [agonist] vs. response, variable slope using GraphPad Prism 9. Each condition was analyzed in triplicate, with four independent replications.

### Coactivator recruitment assay

Ligand-induced recruitment of SRC-1 coactivator peptide to the ligand binding domain of FXR was assayed by time-resolved FRET using LanthaScreen technology according to the manufacturer’s instructions (Thermo Fisher Scientific, cat# A15140). Concentration of test compounds ranged from 10 nM to 100 μM in 1.0% DMSO. TR-FRET signals were assessed after 5.5 hours of incubation at room temperature, at wavelengths of 490 nm (donor) and 520 nm (acceptor) using a Tecan Spark microplate reader. FRET ratios (acceptor:donor emission) are expressed relative to the ratio of the solvent control (*i*.*e*. 1.0% DMSO = 1.0). Dose-response curves were fitted by non-linear regression [agonist] vs. response, variable slope using GraphPad Prism 9. Each condition was analyzed in quadruplicate, with a single replication.

### Reporter assays

#### cAMP

293T cells, grown in 24 well plates until 70-80% confluency, were transfected (Lipofectamine LTX, Thermo Fisher Scientific) with a mixture of pCMV3-TGR5 (0.35 ng), pGL4-CRE-Fluc (15 ng), pRL-TK (1.5 ng) and pCMV3-GFP (83 ng). After 24 hours, cells were exposed for 5 hours to solvent (0.1% DMSO), positive control (10 μM Forskolin A) or test compounds in serum-free medium, and subsequently lysed by repeated freeze-thawing.

#### FXR

293T cells, grown in 24 well plates until 70-80% confluency, were transfected (Lipofectamine LTX, Thermo Fisher Scientific) with a mixture of pCMV3-SLC10A1/pCMV3-SLC10A2 (0 or 1 ng), pcDNA3.1-FXRα2 or pcDNA3.1-FXRα4 (10 ng), pcDNA3.1-RXRα (2 ng), pGL3-*Shp_e*-FLuc (5 ng), pRL-TK (2 ng) and 81 ng or 80 ng pCMV3-GFP (added to reach 100 ng plasmid per well). 24 hours after transfection, cells were exposed to solvent (0.1% DMSO), positive control (10 μM OCA) or test compounds in serum-free medium for a further 24 hours and lysed.

Firefly and *Renilla* luciferase activity was measured in cell lysates using the Dual-Luciferase Reporter assay system (Promega) in a GloMax navigator luminometer (Promega). Data are expressed as normalized ratios (firefly:*Renilla* signal), relative to the solvent control (*i*.*e*. DMSO=1.0). Conditions were tested in quadruplicate, with 3 independent replications.

### Cytokine release

RAW264.7 cells at 70-80% confluency were pre-treated for 1 hour with solvent (0.1% DMSO), positive control (50 μM TLCA) or the indicated test compounds in serum-free medium. Next, an optimized, non-toxic dose of LPS (3 ng/mL; from *E. coli* O128:B12, Sigma-Aldrich) was added and incubations continued for 23 hours. Conditioned media were harvested, and supernatants (10 min, 300*g*) were assayed for IL-6 and TNFα content by ELISA (RnD Systems, DY406 & DY410). Cytokine release was normalized to cellular protein mass. Conditions were tested in quadruplicate, and experiments were repeated thrice.

### Quantification of MBSCs

MBSCs were incorporated into an existing LC-MS assay for quantifying bile salts, essentially according to García-Cañaveras *et al*. (25). In brief, plasma and bile (pre-diluted with water) samples were deproteinized with 2 volumes of methanol and 1 volume of a mix of deuterated internal standards in isopropanol. Following vigorous mixing and centrifugation (15 min., 50,000g, 4°C), supernatant was transferred to sealable micro-insert glass vials. 5.0 μL sample was injected onto a reverse phase column (Acquity UPLC BEH Shield RP18 column, 2.1 x 100 mm, 1.7 μm particle size, Waters) equilibrated with 85%A (95% H_2_O, 5% acetonitrile containing 10 mmol/L ammonium acetate) and operated at 50°C. Gradient elution to 100%B (acetonitrile) was performed using an UltimateTM 3000 quaternary UPLC pump (Thermo Scientific) at a flow rate of 0.7 mL/min at 50°C. For detection, a Xevo TQ-S mass spectrometer (Waters) with an electrospray ionization source was employed in negative ion mode. MBSCs were monitored as tyrosine/phenylalanine/leucine daughter ions using transitions to 180, 164 and 130, respectively. GCDCA-D4 was used as internal standard for MBSCs. Unweighted linear regression was used to calculate sample concentrations from a 12-point standard curve spiked in plasma with a low endogenous bile salt content.

### Statistical analysis

A linear mixed model was used to analyze data from replicated experiments, with replications modelled as correlated random effect. For FXR reporter assays, OCA was included as positive control to verify proper functioning of the cellular assay. OCA had a notably larger effect than the other compounds, thus, impacting on variance structure in the statistical approach. Grubb’s test identified OCA as significant outlier (*P*<0.01) in each of the conditions (+/-transporter). Data points from OCA treatment were therefore not included in the linear mixed model analysis, but are still presented graphically. Treatment effects were compared versus the control situation, applying Bonferroni correction for multiple testing. Data are presented as the average (± SD) of the means of independent replications. A p-value ≤ 0.05 was considered statistically significant. Statistical analyses were conducted in SPSS version 29.0 (IBM) and GraphPad Prism 9.0 (GraphPad Software, La Jolla, CA, USA).

## Results

### MBSCs are activating ligands for TGR5

TGR5 is a cell surface G-protein coupled receptor for both unconjugated and conjugated bile salts, displaying highest affinity for secondary species (26). To test whether MBSCs act as ligands of TGR5, we first assessed receptor activation via recruitment of a mini G_s_ protein, using a proximity-based bioluminescence resonance energy transfer assay (23,24). TLCA, the most potent endogenous TGR5 ligand known thus far, elicits concentration-dependent recruitment of a mini-G_s_ protein, with an EC_50_ value (1.7 μM) similar to reported values (27). (**Fig. 1A**). Each of the CDCA-based MBSCs activated TGR5 with similar efficacy as TLCA and host-derived glycine/taurine-conjugates of CDCA, albeit with lower potency (in order of decreasing affinity: TLCA > TCDCA=GCDCA=CDCA > LeuCDCA=PheCDCA > *D*-TyrCDCA=*L*-TyrCDCA, **Fig. 1A, Table 1**). CA-based MBSCs were weaker TGR5 agonists compared to their CDCA-based counterparts, with EC_50_ values approximately 7-fold higher (potency order: TLCA > TCA=PheCA=GCA > CA=*D*-TyrCA > LeuCA, **Fig. 1A, Table 1**).

**Table 1.**
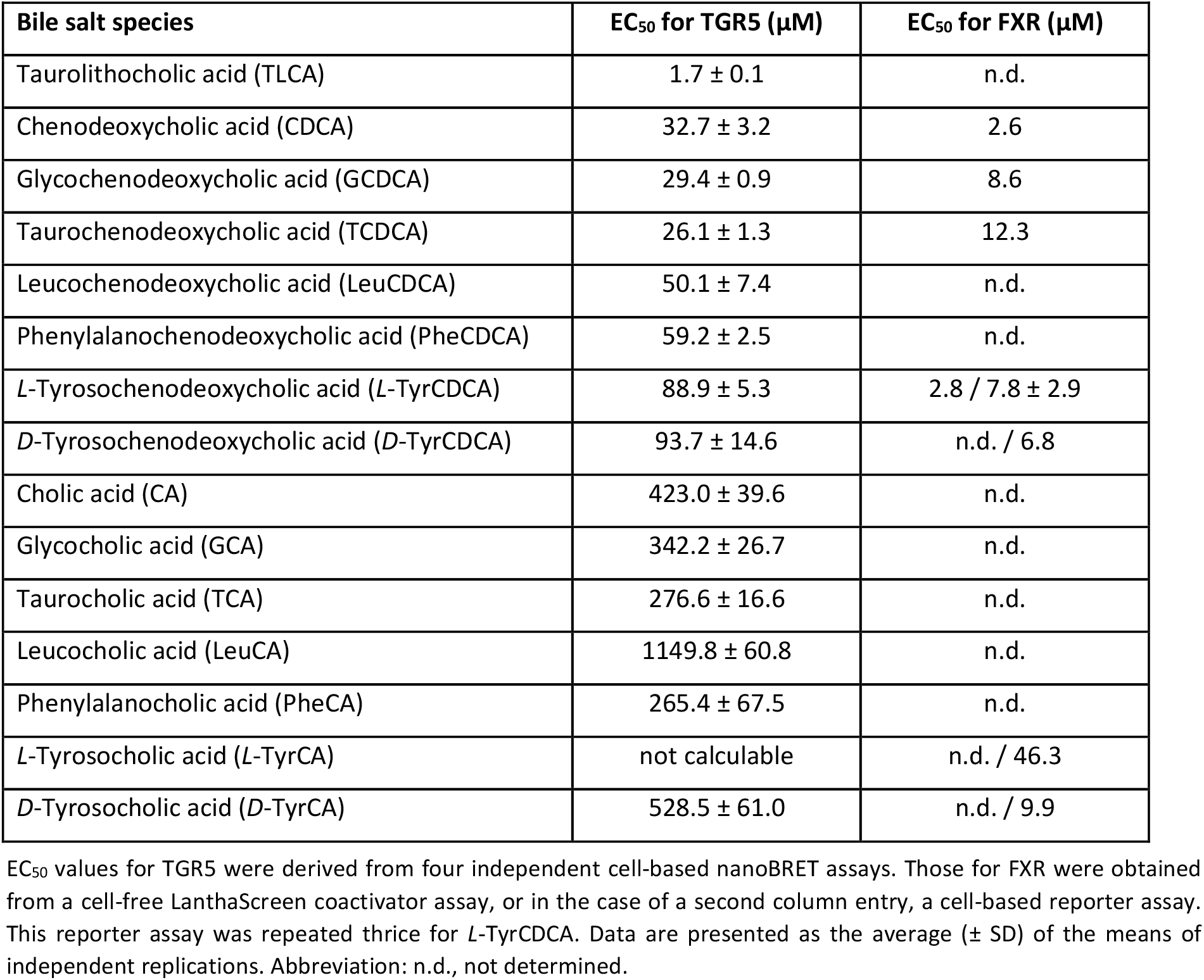
Overview of affinity constants of microbial bile salt conjugates for the main host bile salt receptors.

| **Bile salt species** | **EC<sub>50</sub> for TGR5 (μM)** | **EC<sub>50</sub> for FXR (μM)** |
| --- | --- | --- |
| Taurolithocholic acid (TLCA) | 1.7 ± 0.1 | n.d. |
| Chenodeoxycholic acid (CDCA) | 32.7 ± 3.2 | 2.6 |
| Glychenodeoxycholic acid (GCDCA) | 29.4 ± 0.9 | 8.6 |
| Taurochenodeoxycholic acid (TCDCA) | 26.1 ± 1.3 | 12.3 |
| Leucochenodeoxycholic acid (LeuCDCA) | 50.1 ± 7.4 | n.d. |
| Phenylalanochenodeoxycholic acid (PheCDCA) | 59.2 ± 2.5 | n.d. |
| *L*-Tyrosochenodeoxycholic acid (*L*-TyrCDCA) | 88.9 ± 5.3 | 2.8 / 7.8 ± 2.9 |
| *D*-Tyrosochenodeoxycholic acid (*D*-TyrCDCA) | 93.7 ± 14.6 | n.d. / 6.8 |
| Cholic acid (CA) | 423.0 ± 39.6 | n.d. |
| Glycocholic acid (GCA) | 342.2 ± 26.7 | n.d. |
| Taurocholic acid (TCA) | 276.6 ± 16.6 | n.d. |
| Leucocholic acid (LeuCA) | 1149.8 ± 60.8 | n.d. |
| Phenylalanocholic acid (PheCA) | 265.4 ± 67.5 | n.d. |
| *L*-Tyrosocholic acid (*L*-TyrCA) | not calculable | n.d. / 46.3 |
| *D*-Tyrosocholic acid (*D*-TyrCA) | 528.5 ± 61.0 | n.d. / 9.9 |

EC<sub>50</sub> values for TGR5 were derived from four independent cell-based nanoBRET assays. Those for FXR were obtained from a cell-free Lanthascreen coactivator assay, or in the case of a second column entry, a cell-based reporter assay. This reporter assay was repeated thrice for *L*-TyrCDCA. Data are presented as the average (± SD) of the means of independent replications. Abbreviation: n.d., not determined.

**Figure 1.**
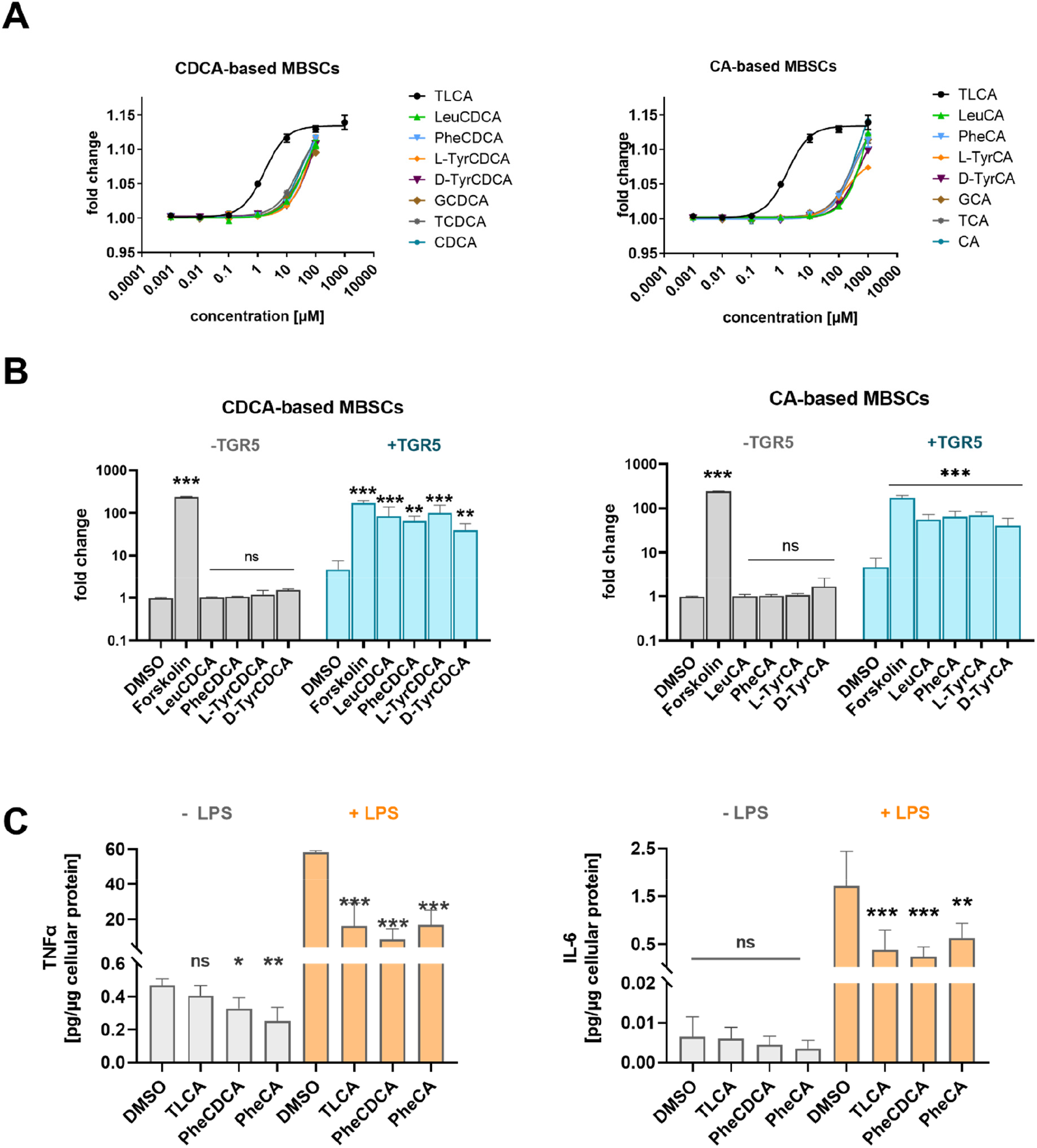
TGR5 activating potential of microbial bile salt conjugates. (**A**) Recruitment of a mini-G_s_ protein to ligand-activated TGR5 was studied in 293 cells expressing tagged protein versions, allowing their interaction to be monitored by bioluminescence resonance energy transfer. TGR5 activation was evaluated for both CDCA-(left panel) and CA-based MBSCs (right panel). Experiments were replicated four times, with three technical replicates per condition in each experiment. (**B**) Signaling downstream of TGR5 was assessed in 293T cells carrying a cAMP-driven luciferase reporter, in the absence (-TGR5) or presence of TGR5 overexpression (+TGR5). TGR5 activation was evaluated for both CDCA-(left panel) and CA-based MBSCs (right panel) at 100 μM. Experiments were repeated three times, with four technical replicates per condition in each replication. (**C**) Inhibition of NF-*κ*B signaling was tested in RAW264.7 cells treated with the indicated test compounds for 1 hour prior to exposure to solvent (-LPS) or 3 ng/mL LPS (+LPS). Cytokine levels in the medium were normalized to cellular protein content. Experiments were replicated twice, with four technical replicates per condition in each test. Data are presented as the average ± SD of independent replications and were statistically evaluated using a linear mixed model. Treatment effects were compared versus the control situation, applying Bonferroni correction for multiple testing. Significance is depicted as * (*P*<0.05), ** (*P*<0.01) and *** (*P*<0.001). **Abbreviations**: CA, cholic acid; CDCA, chenodeoxycholic acid; DMSO, dimethyl sulfoxide; G(CD)CA, glyco(chenodeoxy)cholic acid; IL-6, interleukin 6; LPS, lipopolysaccharide; MBSCs, microbial bile salt conjugates; ns, not significant; T(CD)CA, tauro(chenodeoxy)cholic acid; TLCA, taurolitocholic acid; TNFα, tumor necrosis factor α.

Downstream consequences of TGR5 activation were next evaluated using a reporter assay for cAMP. Effectiveness of the cellular assay was confirmed by incubation with the adenylate cyclase stimulator forskolin, which activated the reporter independent of TGR5 overexpression (**Fig. 1B**). MBSCs elicited strong activation of the reporter in a strict TGR5-dependent manner. Differences in effectiveness among CDCA- and CA-based variants were not apparent at the test concentration (100 μM).

In macrophages, TGR5 activation inhibits nuclear translocation of NF-*κ*B and, consequentially, has anti-inflammatory effects (28). NF-*κ*B pathway antagonism by MBSCs was assessed in RAW264.7 macrophages, which endogenously express TGR5 (29). Activation of NF-*κ*B signaling through the LPS-TLR4 route, resulted in strongly enhanced medium levels of the proinflammatory mediators TNFα and IL-6 (**Fig. 1C**). LPS-induced cytokine level was diminished (TNFα: −3.6 fold, IL-6: −4.6 fold) in cells pre-treated with TLCA, in line with the reported anti-inflammatory action of TGR5 (28,30). Likewise, prior exposure to PheCDCA and PheCA (the most hydrophobic CDCA- and CA-based MBSCs in this study), counteracted LPS-enhanced medium levels of TNFα and IL-6 (**Fig. 1C**). Note that the tested MBSCs somewhat reduced medium TNFα levels in the absence of LPS, which may relate to slight effects on cellular viability (**Fig. S1**).

### MBSCs are substrates for ASBT and NTCP and activate FXR

To assess if MBSCs are FXR agonists, we first determined the effect of *L*-TyrCDCA in a cell-free system to bypass the potential requirement for an uptake transporter. Using a time-resolved FRET assay, we observed concentration-dependent recruitment of SRC1 coactivator peptide to the ligand-binding domain of FXR upon incubation with *L*-TyrCDCA **(Fig. 2A**). The affinity for FXR was similar for *L*-TyrCDCA (EC_50_ = 2.8 μM) and its unconjugated parent bile salt CDCA (EC_50_ = 2.6 μM), and somewhat higher relative to host-conjugated CDCA variants (GCDCA: EC_50_ = 8.6 μM, TCDCA: 12.3 μM) (**Fig. 2A, Table 1**). Affinity of Tyr-MBSCs for FXRα2 was also studied in reporter cells expressing an uptake transporter, revealing composite EC_50_ values of 6.8 μM and 7.8 μM for *D-* and *L*-TyrCDCA, respectively (**Fig. 2B**). A sixfold lower affinity was noted for *L*-TyrCA (composite EC_50_ = 46.3 μM), while the corresponding *D* stereoisomer had affinity (composite EC_50_=9.9 μM) for FXR α2 akin to *D/L*-TyrCDCA (**Fig. S2**).

**Figure 2.**
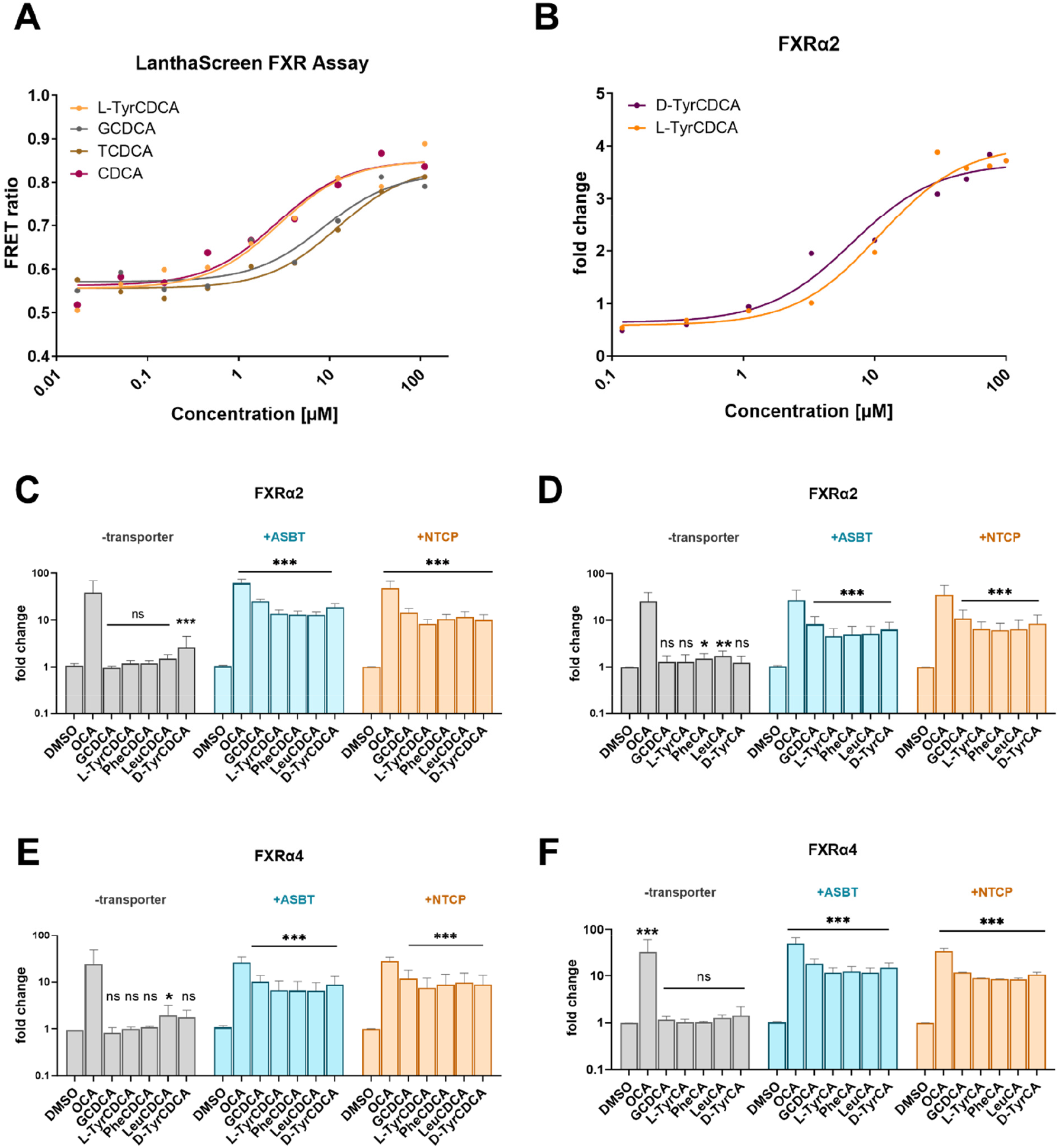
FXR activating potential of microbial bile salt conjugates. (**A**) Ligand-induced recruitment of SRC1 coactivator peptide to the ligand-binding domain of FXR was studied using a TR-FRET based, cell-free assay. Test compounds were evaluated at concentrations ranging from 10 nM to 100 μM, with four technical replicates per condition. (**B**) Affinity of *D-* and *L*-TyrCDCA to activate FXRα2 was studied in 293T cells expressing ASBT and an FXR responsive element-driven luciferase reporter. Transiently transfected cells were exposed to various doses of *L*-TyrCDCA for 16-18 hrs, with four technical replicates per condition. (**C-F**) Activation of FXRα2 and FXRα4 isoforms by MBSCs was investigated using a cell-based reporter assay. Hereto, 293T cells were transiently transfected with the indicated FXRα isoform, in the absence or presence of a bile salt importer (ASBT, NTCP), and incubated overnight with 50 μM CDCA-based MBSCs (panels C, E) or their CA-based equivalents (panels D,F). All conditions were tested in quadruplicate, with 3 independent replications. A linear mixed model was used to analyze data from replicated experiments. Treatment effects were compared versus the control situation, applying Bonferroni correction for multiple testing. Significance is depicted as * (*P*<0.05), ** (*P*<0.01) and *** (*P*<0.001). **Abbreviations**: ASBT, apical sodium-dependent bile acid transporter; DMSO, dimethyl sulfoxide; GCDCA, glycochenodeoxycholic acid; NTCP, Na^+^ taurocholate co-transporting polypeptide; ns, not significant; OCA, obeticholic acid.

FXR activation by each of the 8 MBSC variants available for this study, was further evaluated using a cell-based reporter approach. To this end, 293T cells, which have negligible endogenous *ASBT* and *NTCP* mRNA expression, were transiently transfected with the liver-(FXRα2) or intestinal-enriched (FXRα4) isoform of FXR, in the absence or presence of a sodium-dependent bile salt uptake transporter (**Fig. 2C-F**) (22,31). The membrane-permeable FXR agonist obeticholic acid (OCA) strongly activated the FXR reporter in cells overexpressing FXRα2 or FXRα4, independent of the presence of a bile salt importer. In contrast, notable reporter activation by MBSCs required exogenous expression of either the intestinal (ASBT) or hepatic (NTCP) bile salt uptake transporter. All CDCA-based and CA-based MBSCs were able to activate both FXRα2 and FXRα4 isoforms in the presence of ASBT or NTCP (**Fig. 2CE**). The magnitudes of the effect of individual MBSCs on FXRα2 or FXRα4 were similar at the tested concentration (**Fig. 2C-F**). In contrast with the findings of Quinn *et al*. (12), FXR reporter activity in 293T cells required exogenous FXR expression (**Fig. S3**).

### No substantial entry of MBSCs in the human circulation

To assess if MBSCs enter the host circulation, mesenteric venous and portal blood obtained during abdominal pancreatic surgery was assayed. Demographics and peri-operative serum biochemistry of studied patients are presented in **Table S1**. Despite sensitivity of the LC-MS assay in the low nanomolar range (limit of detection: 0.31-0.63 nmol/L), one or more individual MBSCs could be detected in only 6 out of 30 blood samples (**Table 2**). Levels were above the limit of quantification in a single (cholestatic) patient only, revealing low levels of LeuCA (2.5 nmol/L) and LeuCDCA (1.7 nmol/L) in the venous output from the colon. Note that unconjugated and host-conjugated bile salt species could be quantified in all samples and totaled to a median of 15-36 μmol/L, depending on originating blood vessel.

**Table 2.** Levels of individual microbial bile salt conjugates in human biosamples.

| Study | Sample type | LeuCA<br>(nmol/L) | LeuCDCA<br>(nmol/L) | PheCA<br>(nmol/L) | PheCDCA<br>(nmol/L) | TyrCA<br>(nmol/L) | TyrCDCA<br>(nmol/L) | Total Bile Salts |
| --- | --- | --- | --- | --- | --- | --- | --- | --- |
| **FOCUS** | SMV<br>(n=10) | 1/1<br>[3.4] | 0/0 | 0/0 | 0/0 | 1/0 | 0/0 | 35.9 µmol/L<br>[5.8-266] |
|  | IMV<br>(n=10) | 1/1<br>[2.5] | 1/1<br>[1.7] | 0/0 | 1/0 | 0/0 | 0/0 | 15.3 µmol/L<br>[3.1-220] |
|  | PV<br>(n=10) | 0/0 | 0/0 | 0/0 | 1/0 | 0/0 | 0/0 | 28.6 µmol/L<br>[5.6-216] |
|  | bile<br>(n=14) | 7/7<br>498<br>[71-2163] | 10/9<br>411<br>[65-5111] | 4/3<br>1061<br>[473-2628] | 8/5<br>464<br>[88-1327] | 1/0 | 0/0 | 33.8 mmol/L<br>[2.3-370] |
| **RESCUE** | CV<br>(n=8) | 0/0 | 0/0 | 0/0 | 0/0 | 0/0 | 0/0 | 1.5 µmol/L<br>[0.4-3.4] |
|  | Jejunal<br>chyme<br>(n=8) | 8/8<br>10.2<br>[1.1-56.4] | 8/7<br>6.5<br>[0.8-31.0] | 8/8<br>39.8<br>[3.9-101.3] | 8/7<br>13.1<br>[1.6-56.4] | 5/8<br>4.4<br>[2.6-7.0] | 6/8<br>6.0<br>[2.9-11.9] | 4.4 mmol/L<br>[0.8-7.3] |

The first line in data entries in columns of individual MBSCs states the number of samples in which the particular MBSC could be detected and quantified, respectively. Limit of detection was 0.31 nmol/L for Leu-based MBSCs and 0.63 nmol/L for the other MBSCs. Limit of quantification was 0.63 nmol/L for Leu-based MBSCs and 1.25 nmol/L for the other MBSCs. If quantification was feasible, the second line in data entries provides absolute values, presented as median and -if applicable- [range] (third line). Note that total bile salt levels in plasma and bile/chyme are in micromolar and millimolar units, respectively, and are depicted as median [range]. Abbreviations: SMV, superior mesenteric vein; IMV, inferior mesenteric vein; PV, portal vein; CV, cubital vein.

Unexpectedly, we noted that MBSCs were present in bile of these patients. Leu-based MBSCs could be quantified in over half of the bile samples, with -in that case-median levels of 498 nmol/L (LeuCA) and 411 nmol/L (LeuCDCA) (**Table 2**). Phe-based MBSCs were detected somewhat less frequent, and were quantifiable in 21% (PheCA, median 1061 nmol/mL) and 36% (PheCDCA, median 464 nmol/mL) of the instances. Generally, MBSC variants of Tyr were not present in bile. There was considerable inter-individual variation in biliary MBSC levels that remained when expressed relative to the total amount of bile salts. In samples with quantifiable levels, total MBSCs comprised a minor fraction (14.9 [1.5-213] ppm) of total bile salts in bile.

MBSCs were assayed in a second study, involving patients with acute intestinal failure and verified occurrence of MBSCs in their small intestinal lumen (18, 21). Despite examining those patients with the highest MBSC levels in their chyme, none of the individual MBSCs could be detected in corresponding systemic venous blood (**Table 2**).

## Discussion

In this study, we examined whether microbially conjugated bile salts enter the human circulation and can activate the main host bile salt receptors at the cell surface (TGR5) and in the nucleus (FXR). The key findings are that the tested MBSCs are substrates for the bile salt uptake transporters in the intestine (ASBT) and the liver (NTCP) and are activating ligands of TGR5 and FXR *in vitro*. Despite potential for intestinal absorption, the studied MBSCs were generally not detected in the venous output from the small and large intestines in the studied patients. Surprisingly, specific MBSCs were readily detected in human bile, where they comprised a minor fraction of total bile salts. Given their low systemic levels, surplus of other bile salt species with similar affinities for these receptors, the studied MBSCs are unlikely to have an impact on enterohepatic TGR5/FXR signaling in the host.

*N*-amidates of cholic acid with non-canonical amino acids (*viz*. Leu, Phe and Tyr) were first reported three years ago and shown to be of gut microbial origin, with two *Enterocloster boltae* strains capable of producing them *in vitro* (12). The number of MBSCs (also referred to as microbially conjugated bile acids (14) and bacterial bile acid amidates (32)) has vastly expanded since, with variation in parent bile salt and linked amino acid (both proteinaceous and non-proteinaceous, as well as dipeptides) giving rise to recognition of over hundred species nowadays (15,16). This was paralleled by identification of additional bacterial producers, with MBSC formation appearing a common feature of gut bacteria. Previously unrecognized amine *N*-acyl transferase activity of bile salt hydrolases (bsh), a ubiquitous bacterial enzyme that catalyzes deconjugation of host bile salts, appears to be involved in formation of MBSCs (33). Additional biosynthetic pathways likely exist, as suggested by MBSC production by bacteria seemingly devoid of *bsh* (33). Notwithstanding the advances in basic aspects of MBSCs, insight into their biological function(s) is limited. To appreciate a role in host bile salt signaling, we studied interaction of MBSCs with the key bile salt receptors TGR5 and FXR. MBSCs are not commercially available and were prepared by chemical synthesis. Studied MBSCs encompassed the three initially discovered MBSCs, their CDCA-based equivalents, and the respective conjugates with the *D* enantiomer of tyrosine. Previously, we showed that the latter are resistant to cleavage by bacterial bsh and pancreatic carboxypeptidases (18).

MBSCs were found to activate the G protein-coupled receptor TGR5, as evidenced by recruitment of stimulatory G_s_ subunit surrogate (mini-G_s_ protein) and enhanced expression of a cAMP-driven reporter (**Fig. 1AB**). CDCA-based MBSCs were generally more potent TGR5 agonists compared to their CA-based equivalents **(Table 1)**, in line with the reported higher affinity of this receptor for hydrophobic and secondary bile salts (28). This was also reflected in the potency of individual CDCA-based MBSCs (in order of decreasing affinity: LeuCDCA=PheCDCA > *D*-TyrCDCA=*L*-TyrCDCA) and their elution order in reverse phase chromatography (TyrCDCA: R_t_ 6.89 min, LeuCDCA: R_t_ 8.20 min, PheCDCA: R_t_ 8.70 min). Phe-based MBSCs were further evaluated for their anti-inflammatory potential and found to diminish LPS-induced release of proinflammatory cytokines TNFα and IL-6 (**Fig. 1C**). This is congruent with an NF-*κ*B inhibitory effect of TGR5 activation by these MBSCs (30). The initially discovered MBSCs were reported to activate hepatic FXR in mice (12). Despite repeated oral gavage, MBSCs could not be detected in portal or systemic blood in this study, and it could not be ruled out that the observed repression of FXR target genes was due to cleavage of MBSCs and effects of the cholic acid moiety itself (12). Here, we observed that *L*-TyrCDCA activated FXR, as supported by recruitment of a coactivator peptide to its ligand binding domain in a cell-free system, and activation of a cellular FXR reporter (**Fig. 2AB**). The EC_50_ values (2.8 and 7.5 μM, resp.) were comparable to values for unconjugated and host-conjugated CDCA (**Table 1**) (34,35) Moreover, each of the tested MBSCs, either with CA or CDCA backbone, profoundly activated the cellular FXR reporter, provided that ASBT or NTCP was present (**Fig. 2CD**).

It is plausible that MBSCs must be absorbed in order to initiate signaling via TGR5 or FXR, although there is debate on whether TGR5 in enteroendocrine cells is expressed at the apical membrane and can, thus, be directly activated by luminal bile salts. Our findings support that the tested MBSCs can enter the portal circulation via ASBT, which in the intestines is primarily expressed in the terminal ileum, and may then undergo hepatic extraction by NTCP. Quantitative information on MBSC levels is largely lacking, with limited -but valuable-insights from analysis of fecal matter (36,37), human jejunal chyme (18), and luminal content sampled at various sites along the intestines in mice (12, 14) and human (36,38). Apart from revealing associations with inflammatory bowel disease and cystic fibrosis, the available data indicates that MBSCs are also present in the small intestinal lumen in at least healthy subjects and patients with acute intestinal failure. Hence, it is likely that MBSCs are available as substrates for uptake by ASBT. To the best of our knowledge, we present the first data on occurrence of MBSCs in the human circulation. Using an LC-MS assay with sensitivity in the low nanomolar range, we first examined MBSCs in blood not yet subjected to first-pass hepatic clearance, thus, offering the best prospects for detection of MBSCs. Despite notable total bile salt levels in these vessels, individual MBSCs were generally not detected, let alone at levels that allowed quantification (**Table 2)**. Non-cholestatic patients were included in this initial analysis (**Table S1**), making it unlikely that abrogated intestinal bile inflow common to this patient category, accounted for the negligible MBSC levels in the (portal) circulation. We previously reported that MBSC levels in jejunal chyme of patients with acute intestinal failure, add up to 39 ppm of total bile salts. In matched plasma of “chyme MBSC-positive” patients, individual MBSCs were again not detectable (**Table 2)**. Considering an obligate 4-fold dilution of samples as part of analytical work-up and a limit of detection of 0.31-0.63 nmol/L, individual MBSCs at concentrations of 2.5 nmol/L or higher, can be detected with our assay. Hence, entry of MBSCs into the host circulation appears non-substantial. Cell-surface or intracellular degradation of MBSCs by enterocytic carboxypeptidases could be at play. Given the substrate promiscuity of OSTαβ (39), it is unlikely that MBSCs are not recognized by this transporter and are retained in the enterocyte rather than being released into the mesenteric venules. The apparent lack of MBSCs in the circulation of human subjects is reminiscent of the failure to detect LeuCA and TyrCA in portal blood of mice receiving these MBSCs by gavage (12).

Surprisingly, we detected MBSCs in bile samples from patients undergoing pancreatic surgery (**Table 2**). Leucine-based MBSCs appeared most abundant and were detectable in more than half of the bile samples. The source of biliary MBSCs is unclear. Ascending bacteria or bile duct-colonizing species, introduced by *e*.*g*. endoscopic procedures, may synthesize them from abundant precursor pools in bile. It cannot be excluded, however, that these atypical bile salt conjugates may be produced by the liver. MBSCs comprise up to 213 ppm of total biliary bile salts, and this low abundant fraction may have been overlooked in prior research employing targeted analytics. Anecdotical reports on atypical bile salt conjugates of hepatic origin (*e*.*g*. ornithinocholic acid) are available in the older literature (40). Independent of their origin, the presence of MBSCs in bile raises the question whether these bile salt species recirculate enterohepatically, like host-derived bile salt conjugates. Improved analytical sensitivity will be important to address this point, as bile is presumably a highly concentrated source. In contrast, MBSCs in the bloodstream may simply be overlooked by current analytical limitations.

We demonstrated that affinity of the tested MBSCs for TGR5 and FXR is comparable to, but certainly not higher than, host-derived bile salt conjugates or unconjugated species. Even if efficacy of MBSCs is somewhat higher, trivial levels in the circulation and surplus of equipotent bile salt species, disfavor a major contribution of the tested MBSCs to signaling via TGR5 and FXR. Additional bile salt receptors exist and may be targeted by MBSCs as already shown for PXR (32). Still, above arguments suggest that the primary role of MBSCs may lie outside direct modulation of host signaling.

MBSCs may contribute to inter-microbial communication. Initial tests indicated that MBSCs (*viz*. LeuCA, PheCA, TyrCA), however, did not affect microbial composition *in vitro* (12). Microbial bile salt modification is regarded as a protective mechanism that offers adapted bacterial species an advantage (8). The hallmark reaction in secondary bile salt formation is 7α-dehydroxylation, with few bacterial species harboring the *bai* operon that encodes the relevant enzymatic machinery (41). One example is *Clostridium scindens* which inhibits germination of the pathogen *Clostridium difficile* by producing secondary bile salts (42). Recently, Foley and colleagues reported that certain MBSCs (*viz*. TyrCA, PheCA, PheβMCA) inhibit spore germination of *C. difficile in vivo* (14). Growth of *C. difficile* was not inhibited in the presence of TyrCA and PheCA, but significantly delayed by Tyr- and Phe-variants of β-muricholic acid. Bacterial reconjugation may be a broad mechanism to thwart growth-inhibitory effects of unconjugated bile acids (42).

Our study has several strengths and limitations. Analysis of patient materials allowed a first glimpse into systemic MBSC levels, or apparent lack thereof, in a human context. The enigmatic finding of MBSCs in bile, raises the provoking question whether non-canonical bile salt amidates are exclusively produced by microbes. A limitation of our study is that it is unclear if our selection of 6 MBSCs is representative of the vastly expanding repertoire of MBSCs. Without a commercial source, our choice for chemical synthesis was based on the three initially discovered MBSCs. It can be reasoned that these three were discovered in the first place, because they represented abundant MBSC species, as indeed reflected in later studies (14,37,38). Moreover, the studied MBSCs presumably behave as classical, monovalent bile salt anions. Amidation with acidic or basic amino acids, however, will result in divalent or zwitterionic species, with presumably distinct biophysical (*e*.*g*. membrane permeating) and biological properties. Hence, certain MBSCs may be less susceptible to degradation by host and microbial enzymes and may be absorbed in amounts relevant for host bile salt signaling. Along this line, gavage of mice with *D*-TyrC(DC)A, which is refractory to *in vitro* degradation by pancreatic carboxypeptidases and *bsh* (18), may shed light on enzymatic breakdown as cause of low systemic levels. Note that it is currently unresolved if *D* amino acids are actual substrates for bacterial bile salt reconjugation. Above notions offer exciting prospects for experimental and analytical follow-up efforts.

In conclusion, we demonstrated that the tested MBSCs activate the cell surface bile salt receptor TGR5. Moreover, they are activating ligands of the transcription factor FXR, with a cellular bile salt uptake system required for FXR activation. The studied MBSCs are substrates for both ASBT and NTCP, but their entry into the portal circulation is non-substantial in humans. MBSCs are readily detected in human bile, where they represent a minor portion of total bile salts. Since affinity of the evaluated MBSCs for TGR5 and FXR is not superior to host-derived bile salt conjugates, the trivial levels are unlikely to contribute to host bile salt signaling via these receptors. The biological functions of MBSCs remain to be defined and may be restricted to an ecological role in the gut lumen.

## Abbreviations

ASBT: apical sodium-bile salt transporter
BRET: bioluminescence resonance energy transfer
BSA: bovine albumin serum
*C. difficile*: *Clostridium difficile*
CA: cholic acid
cAMP: cyclic adenosine monophosphate
CDCA: chenodeoxycholic acid
DCA: deoxycholic acid
DMSO: dimethyl sulfoxide
FXR: farnesoid X receptor
GCA: glycocholic acid
GCDCA: glycochenodeoxycholic acid
GLCA: glycolithocholic acid
IL-6: interleukin 6
LCA: lithocholic acid
LC-MS: liquid chromatography–mass spectrometry
LeuCA: leucocholic acid
LeuCDCA: leucochenodeoxycholic acid
LPS: lipopolysaccharide
MBSC: microbial bile salt conjugate
MCA: muricholic acid
NF-*κ*B: nuclear factor-*κ*B
NTCP: Na^+^ taurocholate co-transporting protein
NMR: nuclear magnetic resonance
OCA: obeticholic acid
PheCA: phenylalanocholic acid
PheCDCA: phenylalanochenodeoxycholic acid
TCA: taurocholic acid
TCDCA: taurochenodeoxycholic acid
TGR5: Takeda G protein-coupled receptor 5
TLCA: taurolithocholic acid
TNFα: tumor necrosis factor α
TR-FRET: time-resolved fluorescence resonance energy transfer
TyrCA: tyrosocholic acid
TyrCDCA: tyrosochenodeoxycholic acid.

## Acknowledgements

We thank Prof. Saskia van Mil (Utrecht University Medical Center) for providing us with the pcDNA3.1 vectors containing human FXRα2 and FXRα4, pcDNA3.1-RXRA-Flag, the pGL3-Shp promoter-reporter construct and the control plasmid pcDNA3.1-GFP-HA. Expert advice on statistical analysis of replicated experiments by Dr. Sander van Kuijk (Maastricht University Medical Center) was greatly appreciated. We are grateful to Dr. Athanassios Fragoulis (Institute for Anatomy and Cell Biology, University Hospital Aachen) for his advice on reporter assays.

## Author contributions

Ümran Ay^1^, Martin Leníček^2^, Raphael S. Haider^3,4,5^, Arno Classen^6^, Hans van Eijk^7^, Kiran V.K. Koelfat^1^, Gregory van der Kroft^1^, Ulf. P. Neumann^1,7^, Carsten Hoffmann^3^, Carsten Bolm^6^, Steven W.M. Olde Damink^1,7^ and Frank G. Schaap^1,7*^

Conceptualization: SOD, FGS

Methodology: UA, ML, HVE, FGS

Formal analysis: UA, ML, RSH, HVE, FGS

Resources: AC, KVK, GVDK, UPN, CH, CB

Writing – original draft preparation: UA, FGS

Funding acquisition: SOD, FGS

## Supplemental Information

**Figure S1.**
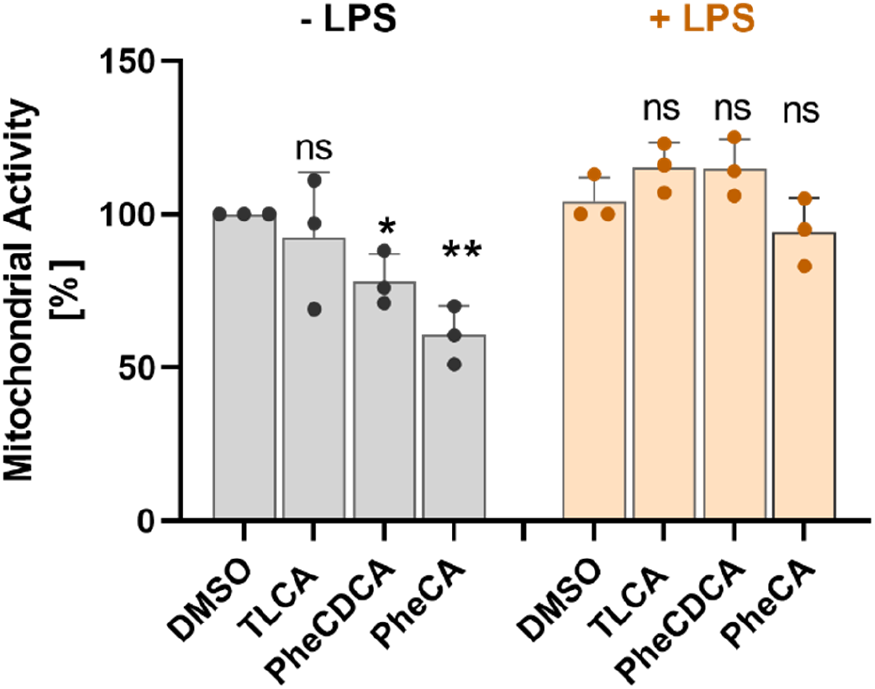
Viability of RAW264.7 macrophages after 1 hour of pre-treatment with the indicated test compounds (0.1% DMSO, 50 μM TLCA, 100 μM PheCDCA and 100 μM PheCA) and 23 hours further incubation without (-LPS) or with (+LPS, 3.0 ng/mL). Mitochondrial activity was taken as measure of cell viability. Data are presented as mean ± SD. All conditions were tested in quadruplicate, with 3 independent replications. A linear mixed model was used to analyze data from replicated experiments. Treatment effects were compared versus the control situation, applying Bonferroni correction for multiple testing. Significance is depicted as * (*P*<0.02) and ** (*P*<0.007).

**Figure S2.**
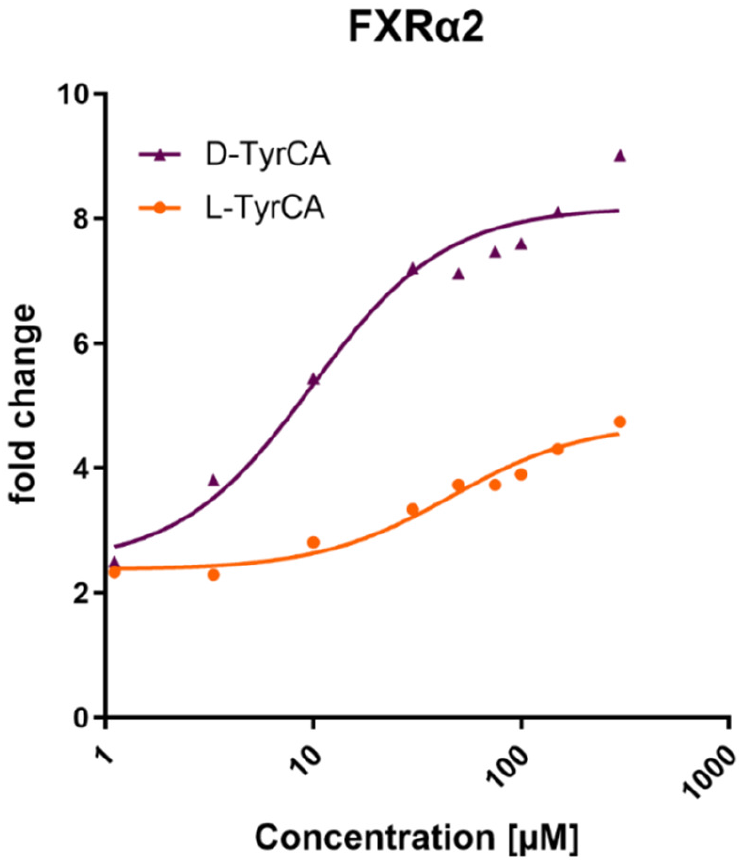
Concentration-dependent activation of an FXR reporter by the *D-* and *L*-stereoisomers of TyrCA. 293T cells were transiently transfected with an expression vector for FXRα2, RXRα, ASBT and an FXR-responsive element driven luciferase reporter. Next, cells were exposed overnight to various concentrations of *D*-TyrCA or *L*-TyrCA, and luciferase activity was determined in cell lysates. All conditions were tested in quadruplicate. Data are presented as mean. Dose-response curves were fitted by non-linear regression [agonist] vs. response, variable slope using GraphPad Prism 9. The *D* stereoisomer (composite EC_50_=9.9 μM) displayed a higher affinity for FXR than *L*-TyrCA (composite EC_50_ =46.3 μM).

**Figure S3.**
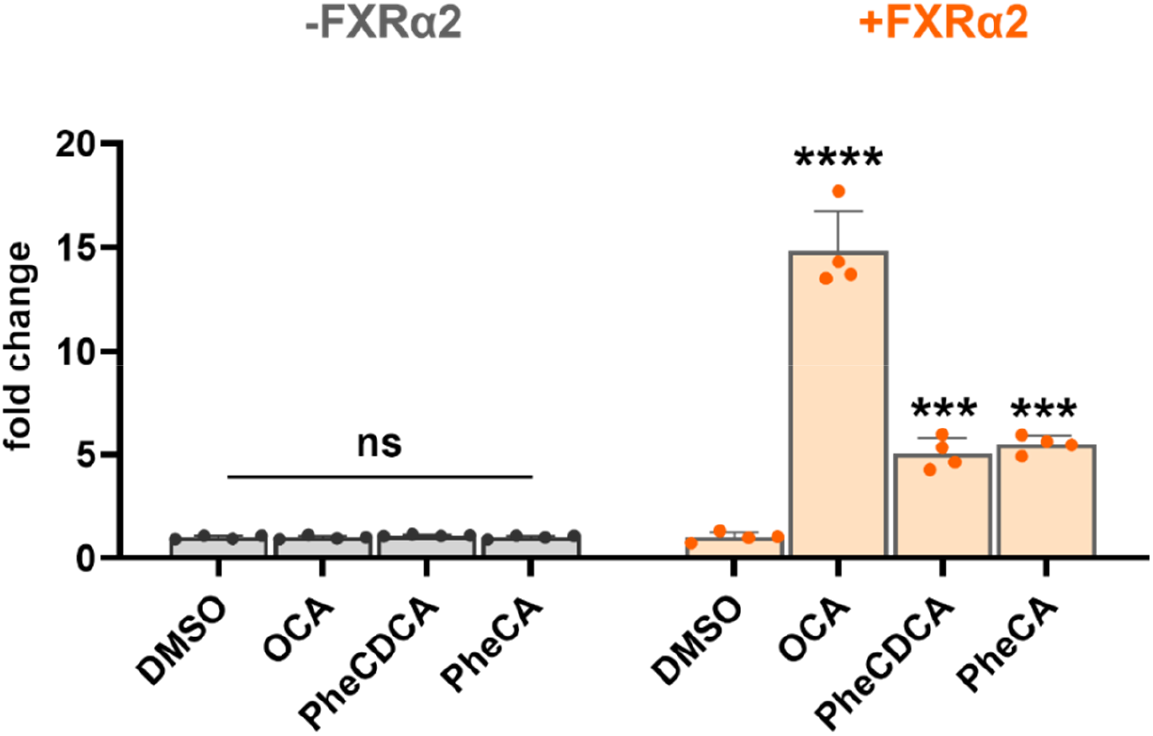
FXR reporter activation requires exogenous FXR expression. 293T cells were transiently transfected with an expression vector for GFP (-FXR, only endogenous *FXR* expression) or FXRα2. All cells were co-transfected with *RXRα, ASBT* and an FXR-responsive element driven luciferase reporter. Cells were then exposed to solvent (0.1% DMSO), the FXR agonist obeticholic acid (OCA, 10 μM), PheCDCA (50 μM) or PheCA (50 μM), and luciferase activity was determined in cell lysates. The luciferase reporter was not activated by OCA and above MBSCs in the absence of exogenous FXR expression. All conditions were tested in quadruplicate. Data are presented as mean ± SD. Effects within FXR groups were evaluated using one-way ANOVA, with Dunnett’s post-hoc test of treatment effects versus the control situation (0.1% DMSO). OCA **** (p<0.0001), PheCDCA *** (0.0006), PheCA *** (0.0002).

**Table S1.** Demographics and pre-operative serum biochemistry of patients undergoing pancreatic surgery.

|  |  |
| --- | --- |
| Gender (female/male) | 3/7 |
| Age (y) | 71 [52-82] |
| BMI (kg/m<sup>2</sup>) | 23 [18-33] |
| Pre-operative biliary drainage (no/yes) | 2/8 |
| Serum bilirubin (μmol/L) | 19.5 [5.4-332] |
| total bilirubin <50 μmol/L (n) | 7 |
| total bilirubin ≥50 μmol/L (n) | 3 |
| ALT (U/L) | 42 [17-396] |
| AST (U/L) | 34 [22-133] |
| ALP (U/L) | 153 [80-554] |
| GGT (U/L) | 129 [39-1785] |
| CRP (mg/L) | 5.2 [0.3-24.3] |

Data are represented as frequency (n) or median [range]. Abbreviations: BMI, body mass index; ALT, alanine aminotransferase; AST, aspartate aminotransferase; ALP, alkaline phosphatase; GGT, gamma-glutamyl transpeptidase; CRP, C-reactive protein.

## References (shared 1^st^ authorships in bold)

1. Hofmann AF. Bile acids: trying to understand their chemistry and biology with the hope of helping patients. Hepatology 2009;49:1403–1418.

2. Arab JP, Karpen SJ, Dawson PA, Arrese M, Trauner M. Bile acids and nonalcoholic fatty liver disease: Molecular insights and therapeutic perspectives. Hepatology 2017;65:350–362.

3. Fickert P, Wagner M. Biliary bile acids in hepatobiliary injury -What is the link? J Hepatol 2017;67:619–631.

4. Fuchs CD, Trauner M. Role of bile acids and their receptors in gastrointestinal and hepatic pathophysiology. Nat Rev Gastroenterol Hepatol 2022;19:432–450.

5. Jansen PL, Ghallab A, Vartak N, Reif R, Schaap FG, Hampe J, Hengstler JG. The ascending pathophysiology of cholestatic liver disease. Hepatology 2017;65:722–738.

6. Albillos A, de Gottardi A, Rescigno M. The gut-liver axis in liver disease: Pathophysiological basis for therapy. J Hepatol 2020;72:558–577.

7. Pabst O, Hornef MW, Schaap FG, Cerovic V, Clavel T, Bruns T. Gut-liver axis: barriers and functional circuits. Nat Rev Gastroenterol Hepatol 2023;20:447–461.

8. Ridlon JM, Kang DJ, Hylemon PB, Bajaj JS. Bile acids and the gut microbiome. Curr Opin Gastroenterol 2014;30:332–338.

9. Campbell C, McKenney PT, Konstantinovsky D, Isaeva OI, Schizas M, Verter J, Mai C, et al. Bacterial metabolism of bile acids promotes generation of peripheral regulatory T cells. Nature 2020;581:475–479.

10. Hang S, Paik D, Yao L, Kim E, Trinath J, Lu J, Ha S, et al. Bile acid metabolites control T(H)17 and T(reg) cell differentiation. Nature 2019;576:143–148.

11. Song X, Sun X, Oh SF, Wu M, Zhang Y, Zheng W, Geva-Zatorsky N, et al. Microbial bile acid metabolites modulate gut RORγ(+) regulatory T cell homeostasis. Nature 2020;577:410–415.

12. Quinn RA, Melnik AV, Vrbanac A, Fu T, Patras KA, Christy MP, Bodai Z, et al. Global chemical effects of the microbiome include new bile-acid conjugations. Nature 2020;579:123–129.

13. Hofmann AF, Hagey LR. Key discoveries in bile acid chemistry and biology and their clinical applications: history of the last eight decades. J Lipid Res 2014;55:1553–1595.

14. Foley MH, Walker ME, Stewart AK, O’Flaherty S, Gentry EC, Patel S, Beaty VV, et al. Bile salt hydrolases shape the bile acid landscape and restrict Clostridioides difficile growth in the murine gut. Nat Microbiol 2023;8:611–628.

15. Lucas LN, Barrett K, Kerby RL, Zhang Q, Cattaneo LE, Stevenson D, Rey FE, et al. Dominant Bacterial Phyla from the Human Gut Show Widespread Ability To Transform and Conjugate Bile Acids. mSystems 2021:e0080521.

16. Wang YZ, Mei PC, Bai PR, An N, He JG, Wang J, Zhu QF, et al. A strategy for screening and identification of new amino acid-conjugated bile acids with high coverage by liquid chromatography-mass spectrometry. Anal Chim Acta 2023;1239:340691.

17. Pedersen KJ, Haange SB, Žížalová K, Viehof A, Clavel T, Leniček M, Engelmann B, et al. Eggerthella lenta DSM 2243 Alleviates Bile Acid Stress Response in Clostridium ramosum and Anaerostipes caccae by Transformation of Bile Acids. Microorganisms 2022;10.

18. Ay Ü Leníček M, Classen A, Olde Damink SWM, Bolm C, Schaap FG. New Kids on the Block: Bile Salt Conjugates of Microbial Origin. Metabolites 2022;12.

19. Guzior DV, Quinn RA. Review: microbial transformations of human bile acids. Microbiome 2021;9:140.

20. Zeng J, Fan J, Zhou H. Bile acid-mediated signaling in cholestatic liver diseases. Cell & Bioscience 2023;13:77.

21. Koelfat KVK, Picot D, Chang X, Desille-Dugast M, van Eijk HM, van Kuijk SMJ, Lenicek M, et al. Chyme Reinfusion Restores the Regulatory Bile Salt-FGF19 Axis in Patients With Intestinal Failure. Hepatology 2021;74:2670–2683.

22. Ramos Pittol JM, Milona A, Morris I, Willemsen ECL, van der Veen SW, Kalkhoven E, van Mil SWC. FXR Isoforms Control Different Metabolic Functions in Liver Cells via Binding to Specific DNA Motifs. Gastroenterology 2020;159:1853–1865.e1810.

23. Leonhardt J, Haider RS, Sponholz C, Leonhardt S, Drube J, Spengler K, Mihaylov D, et al. Circulating Bile Acids in Liver Failure Activate TGR5 and Induce Monocyte Dysfunction. Cell Mol Gastroenterol Hepatol 2021;12:25–40.

24. Wan Q, Okashah N, Inoue A, Nehmé R, Carpenter B, Tate CG, Lambert NA. Mini G protein probes for active G protein-coupled receptors (GPCRs) in live cells. J Biol Chem 2018;293:7466–7473.

25. García-Cañaveras JC, Donato MT, Castell JV, Lahoz A. Targeted profiling of circulating and hepatic bile acids in human, mouse, and rat using a UPLC-MRM-MS-validated method. J Lipid Res 2012;53:2231–2241.

26. Kawamata Y, Fujii R, Hosoya M, Harada M, Yoshida H, Miwa M, Fukusumi S, et al. A G protein-coupled receptor responsive to bile acids. J Biol Chem 2003;278:9435–9440.

27. Nakhi A, McDermott CM, Stoltz KL, John K, Hawkinson JE, Ambrose EA, Khoruts A, et al. 7-Methylation of Chenodeoxycholic Acid Derivatives Yields a Substantial Increase in TGR5 Receptor Potency. J Med Chem 2019;62:6824–6830.

28. Keitel V, Stindt J, Häussinger D. Bile Acid-Activated Receptors: GPBAR1 (TGR5) and Other G Protein-Coupled Receptors. Handb Exp Pharmacol 2019;256:19–49.

29. Pols TW, Nomura M, Harach T, Lo Sasso G, Oosterveer MH, Thomas C, Rizzo G, et al. TGR5 activation inhibits atherosclerosis by reducing macrophage inflammation and lipid loading. Cell Metab 2011;14:747–757.

30. Karababa A, Groos-Sahr K, Albrecht U, Keitel V, Shafigullina A, Görg B, Häussinger D. Ammonia Attenuates LPS-Induced Upregulation of Pro-Inflammatory Cytokine mRNA in Co-Cultured Astrocytes and Microglia. Neurochem Res 2017;42:737–749.

31. Dawson PA, Lan T, Rao A. Bile acid transporters. J Lipid Res 2009;50:2340–2357.

32. Gentry E, Collins S, Panitchpakdi M, et al. A Synthesis-Based Reverse Metabolomics Approach for the Discovery of Chemical Structures from Humans and Animals. In. Preprint at researchsquare; 2021.

33. Rimal B, Collins S, Rocha ER, et al. Bile Acids Are Substrates for Amine N-Acyl Transferase Activity by Bile Salt Hydrolase. In. Preprint at Research Square; 2022.

34. Parks DJ, Blanchard SG, Bledsoe RK, Chandra G, Consler TG, Kliewer SA, Stimmel JB, et al. Bile acids: natural ligands for an orphan nuclear receptor. Science 1999;284:1365–1368.

35. Wang H, Chen J, Hollister K, Sowers LC, Forman BM. Endogenous bile acids are ligands for the nuclear receptor FXR/BAR. Mol Cell 1999;3:543–553.

36. Folz J, Culver RN, Morales JM, Grembi J, Triadafilopoulos G, Relman DA, Huang KC, et al. Human metabolome variation along the upper intestinal tract. Nat Metab 2023;5:777–788.

37. Neugebauer KA, Okros M, Guzior DV, Feiner J, Chargo NJ, Rzepka M, Schilmiller AL, et al. Baat Gene Knockout Alters Post-Natal Development, the Gut Microbiome, and Reveals Unusual Bile Acids in Mice. J Lipid Res 2022;63:100297.

38. Shalon D, Culver RN, Grembi JA, Folz J, Treit PV, Shi H, Rosenberger FA, et al. Profiling the human intestinal environment under physiological conditions. Nature 2023;617:581–591.

39. Suga T, Yamaguchi H, Ogura J, Mano N. Characterization of conjugated and unconjugated bile acid transport via human organic solute transporter α/β. Biochim Biophys Acta Biomembr 2019;1861:1023–1029.

40. Myher JJ, Marai L, Kuksis A, Yousef IM, Fisher MM. Identification of ornithine and arginine conjugates of cholic acid by mass spectrometry. Can J Biochem 1975;53:583–590.

41. Florén CH, Nilsson A. Binding of bile salts to fibre-enriched wheat bran. Hum Nutr Clin Nutr 1982;36:381–390.

42. Begley M, Gahan CG, Hill C. The interaction between bacteria and bile. FEMS Microbiol Rev 2005;29:625–651.

